# Neural encoding of innate preference to gravity-defying motion

**DOI:** 10.1101/2025.11.24.690125

**Authors:** Dmitry Kobylkov, Giorgio Vallortigara

## Abstract

The ability to detect animate objects is a fundamental property of the animal visual system. Among the cues used to infer animacy, gravity provides an important reference for identifying animate motion. Earlier work has demonstrated that upward-moving objects, which violate gravitational constraints, elicit spontaneous attention in newborn humans and domestic chicks. However, the neural mechanisms underlying this innate predisposition remain unclear. We recorded neural activity in the nidopallium of one-week-old domestic chicks as they observed stimuli moving upward or downward. In parallel, we analyzed spontaneous behavioral responses with video-based tracking and high-speed accelerometer data. This analysis revealed a robust attentional bias toward upward-moving, gravity-defying stimuli. We identified neurons in the NCL that encode the direction of motion, most of which responded preferentially to upward movement. Moreover, the population activity of direction-sensitive neurons successfully predicted the spontaneous behavioral response to upward-moving objects.

## Introduction

In an unpredictable natural environment, recognizing animate objects such as prey or predators is a core function of the visual system. Therefore, it is not surprising that the detection of certain animacy features is prewired into the brain. Soon after birth, both human infants and young domestic chicks are spontaneously attracted to animacy cues such as face-like configurations (Johnson et al. 1991, Rosa Salva et al. 2010, 2011) and animate motion (Rosa Salva et al. 2016, Di Giorgio et al., 2016).

To identify animate motion, the visual system can rely on different cues, including coherent movement of body parts (biological motion, Regolin et al. 2000) or spontaneous changes in speed and direction (self-propulsion, Mascalzoni et al. 2010). At the same time, the perception of animate motion in terrestrial organisms is strongly shaped by gravity, which constrains the biomechanics of biological motion and affects expected motion direction (e.g., inanimate objects without external force naturally fall downward). In line with this idea, biological-motion stimuli presented upside down, although retaining low-level movement coherence, become less attractive to newborn human infants (Bardi et al., 2014). In a similar experiment, naïve domestic chicks were not able to distinguish apparent movement direction of the point-light walking hen if it was inverted (Vallortigara and Regolin, 2006). Conversely, in astronauts, exposure to microgravity during spaceflight reduces the inversion effect (Wang et al., 2022).

Given the pivotal role of gravity in animate-motion perception, one fundamental property of animate objects is their ability to move against gravity (i.e., upward). Indeed, humans perceive objects that move against the gravity vector as more animate (Szego and Rutherford 2008).

Similarly, young chicks are spontaneously attracted to upward-moving stimuli. When presented with a choice between upward- and downward-moving discs, chicks approached and spent more time near the upward-moving stimulus (Bliss et al., 2023). At the same time, there was no difference in chicks’ responses to accelerating versus uniformly moving stimuli, indicating that gravity violation alone provides a strong trigger for animacy-related behavior.

While behavioral studies have clearly demonstrated the distinct effect of upward-moving stimuli, the neural correlates of this innate bias remain unknown. The general preference observed by Bliss et al. (2023) is a slow, long-lasting behavioral response that differs significantly from the rapid optokinetic response controlled at the subcortical level (Lapsansky et al. 2025). Therefore, the perception of gravity-defying animate stimuli is likely integrated within a broader animacy-processing network that includes higher-order brain areas. In the avian brain, one of the main candidates for this role is the caudolateral nidopallium (NCL). The NCL, a center for multimodal integration, is involved in a variety of cognitive functions, including object categorization (Pusch et al. 2023). In a recent study, we found that neurons in the NCL of young domestic chicks selectively respond to another crucial animacy cue: face-like configurations (Kobylkov et al., 2024). Moreover, the NCL has been proposed as a telencephalic area involved in global motion integration (Wagener and Nieder, 2016). We therefore hypothesized that the NCL might also be involved in the perception of animacy-related motion cues.

Classical behavioral paradigms used to study innate predispositions are not easily transferable to in vivo electrophysiological experiments. In particular, indirect behavioral measures such as time spent near a preferred object or approach latency are difficult to relate to fast neural responses. Moreover, to identify robust correlations between neural activity and behavior, multiple trial repetitions per subject are required, rather than the one-trial-per-subject design often used in behavioral research.

Therefore, the aims of this study were: (1) to develop a behavioural analysis pipeline capable of reliably detecting rapid spontaneous responses to intrinsically attractive upward-moving objects, and (2) to identify neural correlates of the behavioural bias toward upward-moving objects in young domestic chicks. To minimize the exposure of young chicks to upward-moving stimuli, we first pretrained them to attend to the screen, where only static discs were presented (Fig. 1A). Red discs were rewarded after the stimulus offset, while green discs were unrewarded. This procedure allowed us to dissociate potential reward effects in the subsequent analyses. After the pretraining period, we recorded neural activity and analyzed the spontaneous behavior of chicks as they observed upward- or downward-moving objects (Fig. 1B, Supplementary Movie 1).

**Figure 1.**
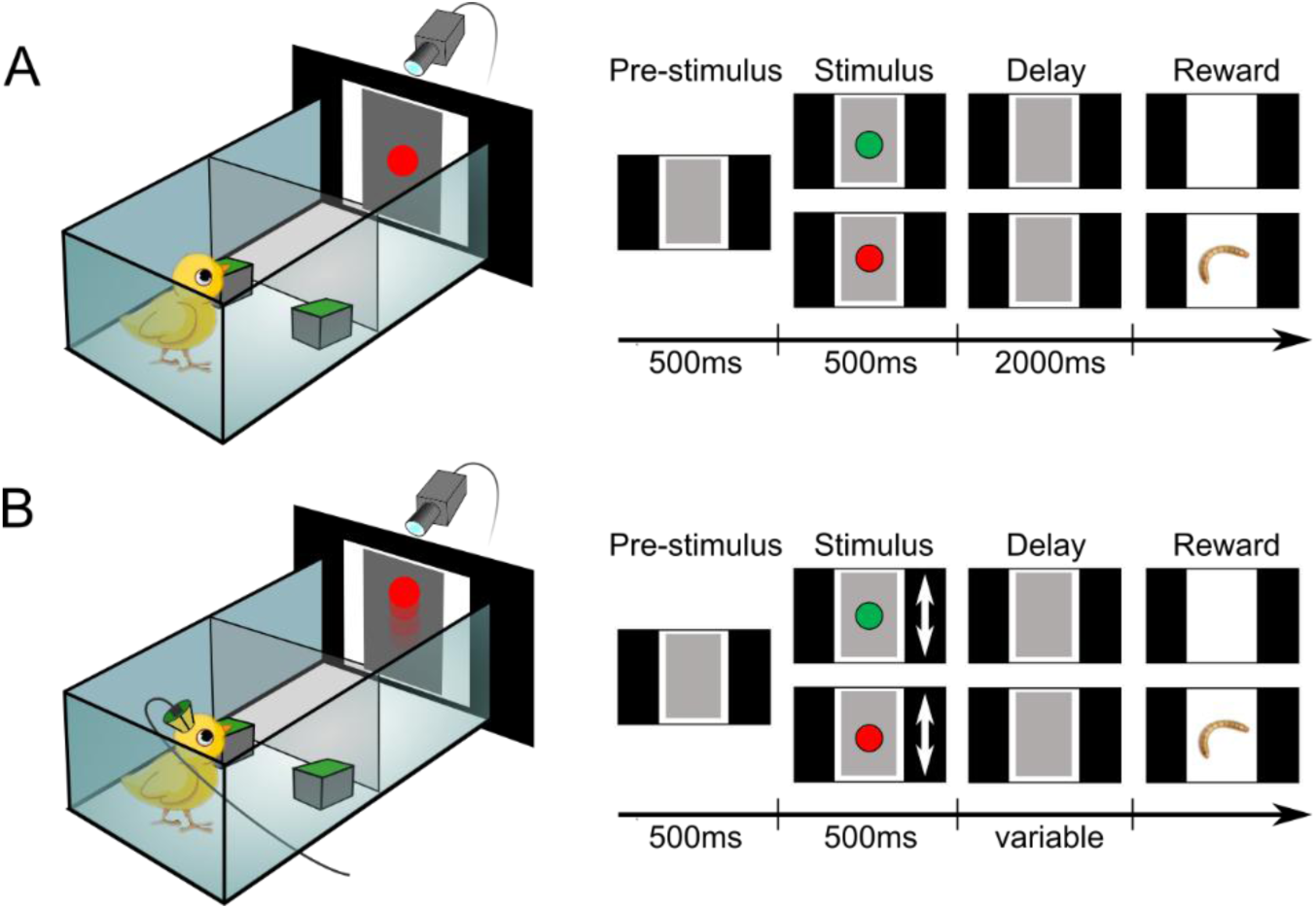
Experimental design. (A) During pretraining phase, animals were presented with static discs. Red discs were subsequently rewarded, while green discs were unrewarded. (B) During the experimental phase, we recorded neural and behavioral response to upward or downward moving stimuli.

## Results

### Spontaneous response to upward moving stimuli

We first analyzed the spontaneous responses of chicks to motion stimuli by tracking the animals using markerless pose estimation with DeepLabCut (Nath et al. 2019). For all subsequent analyses, we included only trials in which animals viewed the stimulus with both eyes or with the contralateral (left) eye. To reveal the posture dynamics during the stimulus presentation, we calculated the average position of the beak (Fig. 2A). We also used the average frame-by-frame displacement of all tracked body parts to quantify overall motion (Fig. 2B).

**Figure 2.**
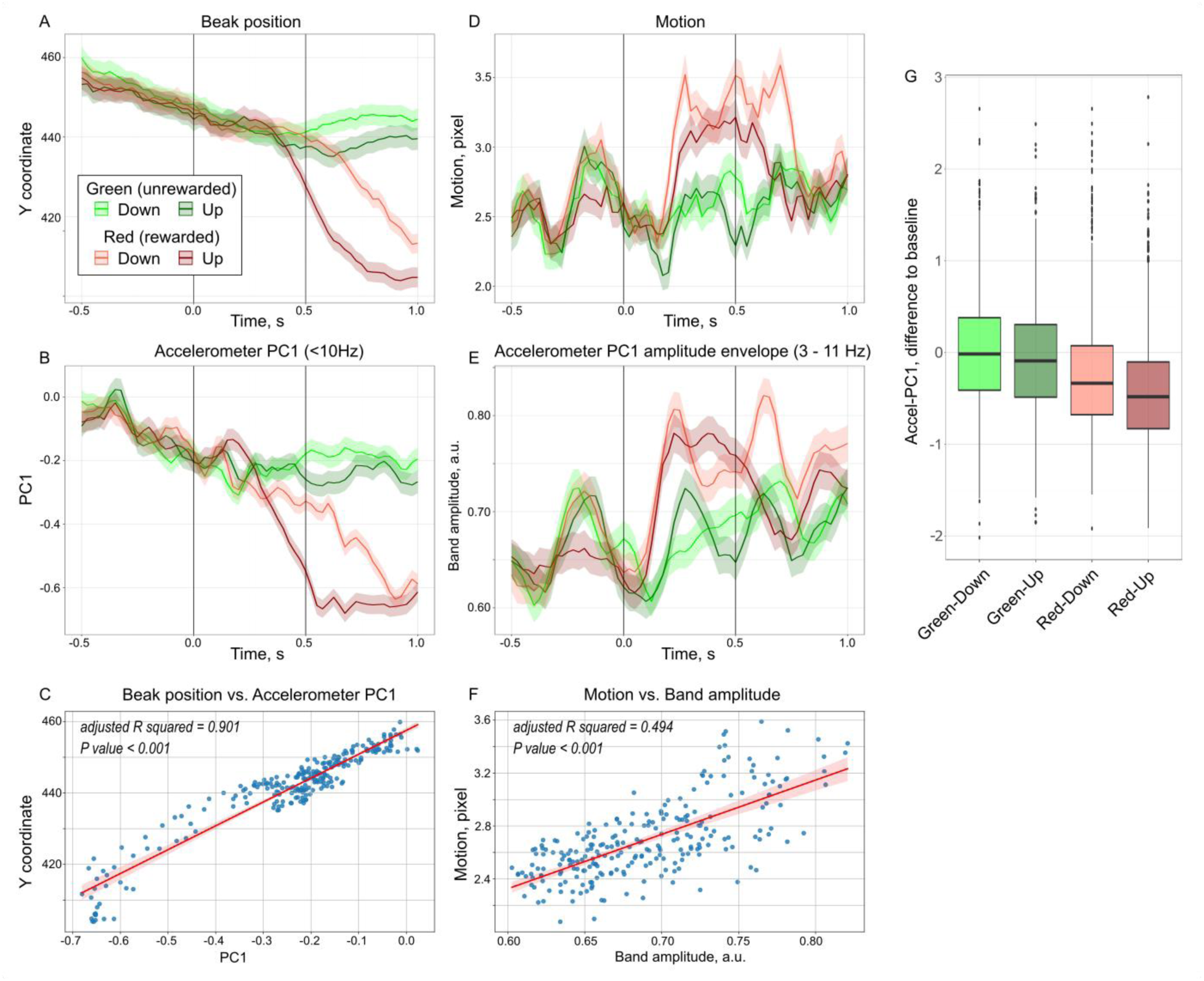
Spontaneous behavioral response to gravity-defying upward-moving stimuli. (A) Vertical average position of the beak estimated from the video analysis with DeepLabCut. (B) Average low-frequency (<10 Hz) component of the accelerometer data (Accel-PC1). (C) Accel-PC1 corresponds well with the beak position and therefore provides a good measure of the posture dynamics. (D) Overall motion response to stimuli based on the video analysis. (E) The amplitude envelope of the Accel-PC1 (in the range 3-11 Hz) corresponds to the overall motion extracted from the video. (F) The amplitude envelope of the Accel-PC1 captures animal overall motion response. (G) Boxplots show distribution of Accel-PC1 difference to the baseline, where negative values correspond to an upward head movement. Importantly, similar to the attention response elicited by the rewarded stimulus, in response to the upward-moving stimulus chicks moved the beak upward (DeepLabCut coordinate system starts at the upper left corner). In all conditions, animals never followed downward-moving stimuli with a corresponding downward head movement. Shaded area in A, B, D, and E correspond to the standard error of the mean (SEM).

In addition to the video analyses, we examined accelerometer data recorded alongside neural activity. To account for slight differences in accelerometer placement across individuals, we performed a principal component analysis on the three axial accelerometer channels. The first principal component (Accel-PC1) was subsequently used as a behavioral measure. We found that the low-frequency (<10 Hz) component of Accel-PC1 corresponded well with posture dynamics captured in the video analysis (Fig. 2C). The average vertical position of the beak was highly correlated with the average Accel-PC1 (Fig. 2E; ordinary least-squares model: p < 0.001, adjusted R² = 0.901). Similarly, the amplitude envelope of Accel-PC1 (3–11 Hz) corresponded to the overall motion extracted from the video (Fig. 2D, F; ordinary least-squares model: p < 0.001, adjusted R² = 0.494). Taken together, the accelerometer data provided reliable measures of both posture and motion at a far higher temporal resolution (30 kHz) than the video tracking (∼25 Hz).

To examine how chicks spontaneously responded to upward-moving stimuli, we analyzed changes in posture (average Accel-PC1) over time across reward conditions (rewarded vs. unrewarded) and movement directions (up vs. down) using a sliding-window two-way ANOVA (100ms window, 10ms steps). We found that posture differed significantly between upward- and downward-moving stimuli beginning 340ms after stimulus onset and lasting for 610ms (Fig. 2B; permutation test: p < 0.001). In contrast, differences in the motion response (amplitude envelope of Accel-PC1) were significant only toward the end of the stimulus, between 470 and 950 ms after onset (Fig. 2E; permutation test: p < 0.001). Because average Accel-PC1 (i.e., posture dynamics) showed faster and more pronounced direction-specific differences, we used this measure for subsequent analyses.

To test whether the response to downward-moving stimuli simply mirrored the response to upward-moving stimuli, we compared the average Accel-PC1 in the direction-specific window (340-950ms) with the preceding baseline level (average in the window -270 – 340ms) for every stimulus condition (Fig. 3). Both for rewarded and unrewarded stimuli, upward moving stimuli elicited significant upward head motion (paired Wilcoxon signed-rank test with p-values adjusted for multiple comparisons: “Red-Up”: *W* = 70320, *p* <0.001, effect size = 0.6; “Green-Up”: *W* = 170164, *p* <0.001, effect size = 0.13). Surprisingly, however, downward-moving stimuli in rewarded trials also resulted in a significant upward head motion (“Red-Down”: *W* = 125505, *p* <0.001, effect size = 0.41). In contrast, unrewarded downward-moving stimuli did not elicit a significant response (“Green-Down”: *W* = 223625, *p* = 0.749, effect size = 0.02).

**Figure 3.**
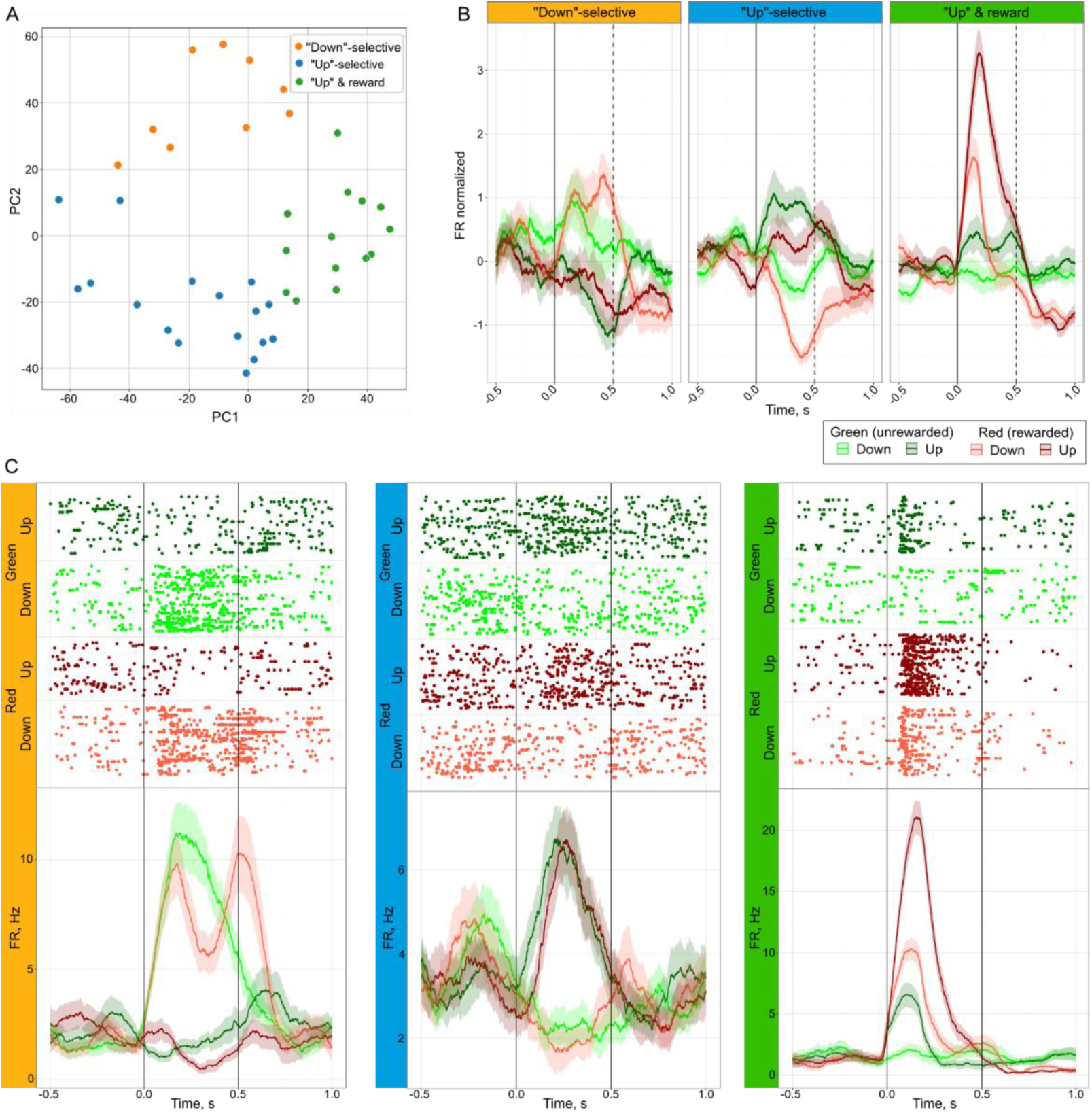
Classification of direction-sensitive neurons. (A) Based on the principal component analysis of neural responses to Upward/Downward and Rewarded/Unrewarded stimuli and k-means clustering three groups of direction-sensitive neurons were identified. (B) Average response of neurons from three clusters. Neural response was gaussian-smoothed (100ms sigma), normalized (z-score) and averaged for each group. (C) Exemplary neurons from each cluster. In the raster plot (top) trials are grouped by stimulus condition and each dot corresponds to a spike. The peristimulus time-histogram (bottom) represents the average Gaussian-smoothed (100ms sigma) neural response to stimuli.

### Neural response to moving stimuli

We recorded neural activity from 213 neurons in the NCL of young domestic chicks while they observed moving stimuli. Using a sliding-window two-way ANOVA (Gaussian-smoothed neural activity: 100ms sigma, 10ms steps), we found that 19% of neurons (N = 40) responded significantly to stimulus direction (Fig. 3).

To further characterize the responses of direction-sensitive neurons, we performed a principal component analysis on their average normalized responses across the four conditions (Rewarded-Up, Rewarded-Down, Unrewarded-Up, Unrewarded-Down). The first two components of the resulting PCA matrix were used to cluster neurons by their response (Fig. 3A). In this way we identified three distinct clusters of neurons (Fig. 3B, C). Only 22.5% of neurons (N = 9) responded more strongly to downward-moving stimuli, whereas the remaining neurons preferred upward motion. The responses of 35% of neurons (N = 14) were modulated not only by direction but also by the reward associated with the stimuli. However, the majority of neurons (42.5%, N=17) were not strongly affected by the reward and responded stronger to upward moving stimuli irrespective of the reward.

### Population response

To determine whether population activity could predict stimulus direction, we performed a time-resolved decoding analysis (Fig. 4A). We trained support vector machines (SVMs) on the firing rate of direction-sensitive neurons in a sliding window of 200ms (20ms steps). Prediction accuracy was above the chance level (estimated by shuffling the training dataset) during the whole stimulus presentation period reaching over 80%. At the same time, SVMs trained on randomly-selected neurons showed overall worse prediction accuracy.

**Figure 4.**
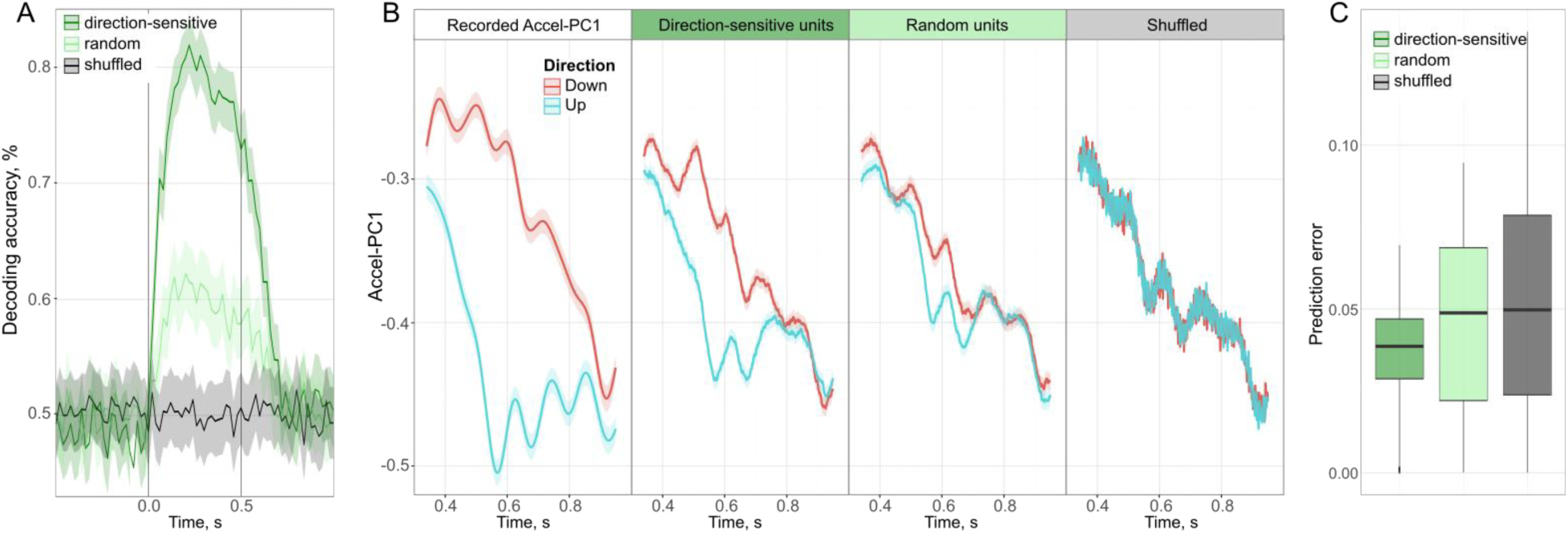
Analyses of the neural population response. (A) Time-resolved decoding accuracy of support vector machines trained on the neural activity of direction-sensitive units (dark-green), randomly-selected units (light-green), or shuffled neural data (black). Shaded area is a 95% confidence interval. (B) Decoding of the observed behavioral response (Accel-PC1) to upward-moving stimuli with a linear regression model trained on activity of direction-sensitive units or randomly-selected units. The chance level was obtained by randomly shuffling the neural data of direction-sensitive units for each time point independently to disturb the direction-dependent but not the time-dependent variability. (C) Prediction accuracy calculated as an absolute prediction error (i.e., lower error values correspond to better decoding).

Furthermore, we tested whether the activity of direction-sensitive neurons could also underlie the behavioral response to the upward-moving stimulus. For this analysis, we trained a linear regression model to decode time-varying behavioral response (Accel-PC1) from the neural activity (Fig. 4B). Using the population activity of direction-sensitive neurons, we successfully decoded the behavioral response within the direction-selective window between 340 and 950ms after the stimulus onset (Fig. 4C; Kruskal–Wallis test: *H* = 179.085, *p* <0.001). Prediction accuracy of this decoder was significantly above the chance level (Pairwise comparisons with permutation test: p<0.001, effect size = 0.26) and better than the decoding performed with randomly selected neurons (p<0.001, effect size = 0.28). Although randomly selected units also performed above chance, the effect size was very small (p<0.001, effect size = 0.06).

## Discussion

So far, studies on innate predispositions have largely relied on simplified behavioral measures of attention, such as time spent near a stimulus or approach rate (Vallortigara et al. 2005, Vallortigara 2021, Bliss et al. 2023). Here, we describe a novel approach for analyzing spontaneous behavioral responses based on both video tracking and high-speed accelerometer data (Accel-PC1). The accelerometer data allowed us not only to analyze the animal’s overall motion but also to capture its posture dynamics.

Analyzing chicks’ posture dynamics, we found that they generally responded to attractive stimuli by moving their heads upward. This was clearly visible when comparing rewarded and unrewarded trials and can therefore be interpreted as an attention response. However, the attention response was significantly modulated not only by reward but also by stimulus direction. Upward head movements in response to rewarded upward-moving stimuli were significantly more pronounced than those elicited by rewarded downward-moving stimuli. Moreover, the unrewarded upward-moving stimulus also elicited a significant attention response. At the same time, it seems unlikely that the response to upward-moving objects reflects simple object-following behavior unconnected to attention. If this were the case, we would expect downward-moving objects to elicit a comparable response – namely, downward head movements. However, even for rewarded downward-moving objects, chicks responded with upward head movements. Moreover, in the unrewarded trials, chicks did not show any significant response to downward motion. Finally, the direction-selective behavioral response persisted for almost 500 ms after the stimulus had disappeared. Together, these findings strongly suggest that upward-moving objects robustly elicit an attentional bias in chicks.

By recording neural responses in the NCL of one-week-old chicks, we found that 19% of neurons significantly modulated their activity according to stimulus motion direction. In line with the observed behavioral bias, the vast majority of direction-sensitive neurons responded more strongly to upward motion. Moreover, cluster analysis revealed that nearly half of the direction-sensitive neurons were influenced primarily by stimulus direction rather than reward. This selective response to motion direction is consistent with previous findings showing that in the NCL of crows performing visual discrimination task 28% of all recorded cells showed direction-selectivity (Wagener and Nieder, 2016). Thus, the NCL appears to be an important brain center for integration of motion cues processed via both the tectofugal (Frost and DiFranco, 1976; Gu et al., 2002) and the thalamofugal visual pathways (Baron et al., 2007).

However, a clear bias to the upward-moving stimuli that we observed in the neural recordings of young chicks contrasts with results from other bird species. In pigeons, Frost and DiFranco (1976) showed that only 7% of motion-selective units in the tectum were tuned to the upward-moving stimuli, while the majority (37%) preferred downward motion. In the entopallium, which gets visual input from the tectum via the nucleus rotundus, the directionality tuning also remains biased towards downward-moving stimuli (Gu et al., 2002). Similarly, the downward bias has been observed in the visual Wulst of awake owls (Baron et al., 2007) and in the NCL of crows (Wagener and Nieder, 2016). One possible explanation for these species-specific differences in direction-selective neural responses may lie in the ecological specialization of chickens. As primarily ground-dwelling animals, chickens may have different directional sensitivities compared to actively flying species. Moreover, in our experiments the chicks were not trained to discriminate motion directions, and we minimized their exposure to upward-moving objects during pretraining by presenting only static stimuli. Therefore, they were far less exposed to upward-moving stimuli than the adult, trained animals used in previous studies (Gu et al., 2002; Wagener and Nieder, 2016). Further experiments in adult chickens are needed to determine whether this effect is experience-dependent or species-specific.

Population analyses of direction-sensitive neurons in the NCL of chicks confirmed that the activity of these neurons encodes the motion direction of the stimulus. Furthermore, by training a linear regression model on the neural responses of direction-sensitive neurons, we were able to successfully decode the animal’s behavioral response. The premotor function of the NCL has been previously described in crows (Kirschhock and Nieder, 2022) and pigeons (Kalt et al., 1999). However, unlike our study, previous work focused on the motor control of voluntary movements. In crows, the sensorimotor neurons were shown to translate perceived numerical stimulus into number of actions (Kirschhock and Nieder, 2022). In pigeons, a small percentage of units in the NCL exhibited higher firing rates prior to beak movements during a Go/NoGo task (Kalt et al., 1999). Instead, we show that NCL activity in chicks predicts spontaneous responses to innately attractive upward-moving stimuli.

In conclusion, our study provides the first insight into the neural mechanisms underlying an innate behavioral bias toward gravity-defying upward-moving objects. By implementing a novel behavioral analysis pipeline combining automated video tracking and high-speed accelerometer data, we characterized spontaneous behavioral responses to upward-moving stimuli. Furthermore, we identified a neural population in the NCL of one-week-old chicks that preferentially responds to upward-moving stimuli and encodes the animals’ posture dynamics during these responses.

## Materials and Methods

### Subjects

We used two domestic chicks (*Gallus gallus domesticus*) from the Aviagen ROSS 308 strain for this study. Fertilized eggs (CRESCENTI Società Agricola S.r.l.–AllevamentoTrepola–cod. Allevamento 127BS105/2) were incubated (Marans P140TU-P210TU) and hatched at 37.7 °C and 60% humidity in darkness in the animal facility. After hatching, chicks were isolated and housed individually in metal cages (28 cm wide × 32 cm high × 40 cm deep) at a constant room temperature of 30–32 °C and a light–dark regime of 14 h light and 10 h dark with food and water provided *ad libitum*. All experimental protocols were approved by the research ethics committee of the University of Trento and by the Italian Ministry of Health (permit number 539/2023-PR).

### Experimental setup

Experiments were performed in a rectangular shaped arena (34 X 54 X 27 cm; W X L X H) with wooden walls and floor covered with non-reflective materials. One wall of the arena was replaced by a computer screen (AOC AGON AG271QG4, 144Hz) used for stimulus presentation (Fig. 1A). The arena was divided in two sections by a metal grid placed 31 cm from the screen. A custom-built automatic reward system consisted of two feeders (left and right) with mealworms, whose lid was attached to a servo motor and controlled by Arduino Uno (Fig. 1A). Stimulus presentation and reward were controlled using Bonsai software (Lopes et al. 2015) with the BonVision toolbox (Lopes et al. 2021). Experiments were recorded with a video camera (Imaging Source DMK 27BUR0135, Germany) at 25 frames per second.

### Training and experimental procedure

On the second day post hatching, chicks learned to peck on mealworms. Between the third and sixth day after hatching, the chicks were habituated to the setup and trained to pay attention to the stimuli (Fig. 1A).

Experimental trials consisted of four stages: 1). To prime attention, the screen turned gray 500ms before stimulus onset; 2). Visual stimuli were presented for 500ms. Red discs were rewarded after the stimulus offset, while green discs are unrewarded. During pretraining phase, the stimuli remain static during the presentation. After electrode implantation (8-11 days after hatching) we presented discs (25mm, 4 degrees angular size) moving upward or downward (angular velocity 66 degrees/s); 3). After stimulus offset and the delay period (variable time between 2000 and 6000ms), one of the feeders was randomly opened for 700-1000ms; 4). The inter-stimulus interval lasted between 2500 and 3000ms.

### Surgery and recordings

On the seventh day after hatching, chicks were fully anaesthetized using Isoflurane inhalation (1.5 – 2.0% gas volume, Vetflurane, 1000mg/g, Virbac, Italy) and placed in the stereotaxic apparatus with a bar fixed at the base of the beak and tilted 45° to ear bars. Local anaesthesia (Emla cream, 2.5% lidocaine + 2.5% prilocaine, AstraZeneka, S.p.A.) was applied to the ears and skull skin before and after the surgery. Metal screws were placed into the skull for grounding and stabilisation of the implant. A small craniotomy was made in the skull above the NCL (1.0 mm anterior to the bregma, 4.5 mm lateral to the midline) on the right. For extracellular recordings, we used self-wired tetrodes made from formvar-insulated nichrome wire (17.78 µm diameter, A-M Systems, USA), which were gold-plated (NeuraLynx, USA) to reduce the impedance to 250 – 350 kOm (controlled by nanoZ, Multi Channel Systems, Germany). Then, a 64-channel drive (Open Ephys, USA) was assembled with 16 tetrodes, implanted, and fixed with quick adhesive silicone (Kwik-Sil, World Precision Instruments, USA) and with dental cement (Henry Schein Krugg Srl, Italy).

After the surgery, the chicks were left to recover until the next day in their home cages. Between the 8th and the 11th day after hatching we recorded neural responses to moving stimuli in the NCL of chicks. After every recording session the tetrodes were manually advanced by approximately 100µm.

Signals were recorded via a low-profile SPI headstage with an integrated 3-axis accelerometer (OEPS-6550, Open Ephys, USA). Acquisition system (Open Ephys) integrated neural data and photodiode input controlling stimulus presentation on the screen. Spike detection and sorting were automatically performed with MountainSort4 (Chung et al. 2017), and all identified units were manually curated using Phy 2.0.

### Data analysis

All statistical analyses and data visualizations were performed in Python using custom-made scripts.

Video recordings were analysed using a markerless pose-estimation technique in DeepLabCut with a model trained on a subset of frames (N=600) to identify “beak”, “left eye”, “right eye”, “left LED” (from headstage), “right LED”, and “headstage base”. Based on the video tracking, we selected only trials in which birds looked with both eyes or with the contralateral (left) eye at the stimulus for at least 50% of the stimulus presentation time. To describe dynamic posture changes we used the average beak position. To analyse the overall motion, we computed average displacement of all tracked body parts. First, for each body part only frames with the high tracking accuracy (likelihood probability >= 0.95) were selected. Then, if two consecutive frames had valid tracking of the same body part, the displacement was calculated as 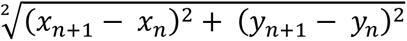, where x and y are coordinates of the body part. Finally, a median of all body parts displacement was calculated.

Accelerometer data were first transformed via principal component analysis using the same transformation matrix for all recordings to eliminate between-individual variation in the accelerometer position. The resulting first principal component (“Accel-PC1”) was used for further behavioural analyses. The low-frequency component of the Accel-PC1 (Butterworth 10Hz low-pass filter of the 4^th^ order) was used as a measure of animal’s posture, while the amplitude envelope of the Accel-PC1 in the frequency range between 3 and 11Hz was used as a measure of animal’s overall motion. We further evaluated how well the accelerometer measures correspond to the parameters extracted from the video by applying a linear regression model (Ordinary Least Squares) and calculating the adjusted R squared.

To reveal the difference in the behavioural response to upward and downward moving stimuli, we performed a sliding-window analysis of variance (two-way ANOVA, 100ms bin window, 10ms step-size) with stimulus condition (rewarded/unrewarded) and the motion direction (up/down) as factors. Accelerometer data (average Accel-PC1 and the amplitude envelope of the Accel-PC1) were first downsampled to 1ms time resolution. Only bins where the response was significantly different (p < 0.01) between upward and downward moving stimuli were selected and the longest consecutive time window was selected. A cluster permutation test was then performed to control for multiple comparisons. For this, all F-values within a significant response window were summed up (F-real) and compared to the sum of F-values resulting from the ANOVA analysis of randomly shuffled trials (F-shuffled). This procedure was repeated 1000 times, and the response window was considered truly significant only if the F-real was higher than 95% of all F-shuffled trials (corresponding to a p < 0.05).

To further quantify the behavioural response to moving stimuli, we compared the average response in the direction-selective window identified earlier with the preceding window of the same length served as a baseline. The difference between baseline and direction-selective window was compared with the Wilcoxon signed-rank test with p-values adjusted for multiple comparisons using the False Discovery Rate (FDR) Benjamini-Hochberg (BH) procedure. An absolute rank-biserial correlation was used as a measure for the effect size.

### Direction-selective neural responses

The neural activity of recorded units was analysed in the 500 ms window starting 100ms after stimulus onset (to account for the visual latency of NCL neurons (Veit et al. 2014) until 100ms after the stimulus offset. First, every trial was smoothed using a Gaussian kernel with 100ms sigma. To identify stimulus-selective neural responses we then performed a sliding-window two-way ANOVA (10 ms bin window, 10 ms step-size) with the stimulus condition (rewarded/unrewarded) and the motion direction (up/down) as factors. Then, only bins where the response was significantly different (p < 0.01) between upward and downward moving stimuli were selected. If a significant response lasted for at least a 100 ms period (10 consecutive bins) we performed an additional cluster permutation test identical to the one used for accelerometer data (described above).

To further characterize direction-selective neurons we classified them into groups based on their response to different stimulus conditions. First, the neural response of every neuron was averaged by stimulus type (Rewarded-Up, Rewarded-Down, Unrewarded-Up, Unrewarded-Down), z-scored, and aligned into one row. Then, the PCA was performed on the matrix with all direction-selective neurons, and the first two components were used to define clusters of neurons (k-means clustering, N clusters = 3).

### Population response

To confirm that neural activity of direction-sensitive neurons indeed encodes the motion direction of the stimulus, we trained support vector machines (SVMs) on the firing rates of direction-sensitive units in the sliding window of 200ms (between 600ms before stimulus onset and 500ms after the stimulus offset, 20ms step). We randomly selected 72 trials per direction (Up/Down) and one trial was randomly assigned for testing. To make the neural activity comparable between units, the firing rates were z-scored using the mean and the SD of the training set only. Then, training and testing of the SVM was performed on every consecutive time bin. For a robust estimation of the decoding accuracy, we performed 1000 iterations of the SVM training and testing, randomly selecting the trials each time. The same procedure was replicated with randomly-selected units that are not direction-sensitive. To estimate the chance level we used shuffled data, where stimulus direction labels were randomly shuffled before training. The confidence interval for each time bin was calculated as 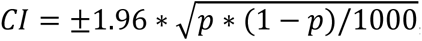, where *p* is the average accuracy for the corresponding time bin.

To reveal the correspondence between the neural activity and the behavioural response to upward-moving stimuli, we performed a decoding analysis using Accel-PC1 data down sampled to 1ms time-resolution as a behavioural output variable. Identical to the SVM analysis, 72 random trials for upward and downward conditions were selected for each unit and one of these trials was randomly assigned for testing. The neural activity was analysed with a rolling average (200ms window, 1ms step) so that each Accel-PC1 value corresponds to the average firing rate in the preceding 200ms time window. Neural data were then z-scored for each unit with the mean and standard deviation estimated for the training trials only. To combine direction-sensitive units from different recordings into one single dataset, the Accel-PC1 response was averaged between recordings and all units were combined as predictors. Finally, the linear regression model was built for each consecutive time point independently and the estimated regression coefficients were used to predict the Accel-PC1 values in the testing trials. The whole procedure was repeated 1000 times to estimate the decoding accuracy. To estimate the chance level accuracy, the correspondence between the neural activity and the Accel-PC1 was disturbed by randomly shuffling the Accel-PC1 values for each time point independently.

The decoding accuracy was evaluated by calculating an absolute error (i.e., absolute difference between average predicted and true values per time bin) and comparing the absolute error distribution between decoders trained with direction-sensitive units, randomly selected units, and shuffled data using a non-parametric Kruskal-Wallis test and a permutation test for post-hoc pairwise comparisons. In addition, an absolute rank-biserial correlation was used as a measure for the effect size.

## Acknowledgements

We would like to thank Mirko Zanon for his comments on the experimental design. We are also grateful to the people at the Animal House Facility for their help with handling the chicks. This study has been supported by funding from the European Research Council under the European Union’s Horizon 2020 research and Innovation Program No. 833504 SPANUMBRA (G.V.), PRIN 2017 ERC-SH4–A 2017PSRHPZ (G.V.), and PRIN 2022 PNRR—Grant Agreement P2022TKY7B (G.V.).

## Author Contributions

D.K. and G.V. designed research; D.K. performed research and analyzed data; D.K. and G.V. wrote the paper.

## Data and code availability

All data and code (Neural recordings and analyses scripts) are available in the main text or in the depository (10.5281/zenodo.17697632) (Kobylkov 2025).

## Supplementary Information

Supplementary Movie 1

Neural activity of an exemplary direction-sensitive unit in the NCL of a young domestic chick shows selective response to the gravity-defying upward-moving stimulus irrespective of the reward. The animal is spontaneously attracted to the upward-moving stimulus, which is visible in the behavioural response (left) and in the corresponding accelerometer data (bottom right). The video recording of the trials corresponds to the neural activity (top right) recorded from one electrode. The neural data are filtered between 300 and 6000Hz. The corresponding visual stimuli are shown in the centre. The recording is slowed down to the quarter of the actual speed.

